# Effects of L-DOPA on gene expression in the frontal cortex of rats with unilateral lesion of midbrain dopaminergic neurons

**DOI:** 10.1101/2020.04.28.063347

**Authors:** Anna Radlicka, Kinga Kamińska, Malgorzata Borczyk, Marcin Piechota, Michał Korostyński, Joanna Pera, Elżbieta Lorenc-Koci, Jan Rodriguez Parkitna

## Abstract

The development of Parkinson’s disease (PD) causes dysfunction of the frontal cortex, which contributes to hallmark motor symptoms and is regarded as one of the primary causes of the affective and cognitive impairments observed in PD. Treatment with L-DOPA alleviates motor symptoms but has mixed efficacy in restoring normal cognitive functions, which is further complicated by the psychoactive effects of the drug. In this study, we investigated how L-DOPA affects gene expression in the frontal cortex in an animal model of unilateral PD. We performed an RNA-seq analysis of gene expression in the frontal cortex of rats with 6-hydroxydopamine (6-OHDA)-induced unilateral dopaminergic lesion that were treated with L-3,4-dihydroxyphenylalanine (L-DOPA), for 2 weeks. We used analysis of variance to identify differentially expressed genes and found 48 genes with significantly altered transcript abundance after L-DOPA treatment. We also performed a weighted gene coexpression network analysis (WGCNA), which resulted in the detection of 5 modules consisting of genes with similar expression patterns. The analyses led to three primary observations. First, the changes in gene expression induced by L-DOPA were bilateral, although only one hemisphere was lesioned. Second, the changes were not restricted to neurons but also appeared to emerge in immune or endothelial cells. Finally, comparisons with databases of drug-induced gene expression signatures revealed multiple nonspecific effects, which indicates that a part of the observed response is a common pattern activated by multiple types of pharmaceuticals in different target tissues. Taken together, our results identify cellular mechanisms in the frontal cortex that are involved in the response to L-DOPA treatment.

## Introduction

Parkinson’s disease (PD) is a neurodegenerative disorder that leads to a progressive loss of dopaminergic neurons of the substantia nigra in the ventral midbrain (Hornykiewicz, 1998). The primary symptoms of PD are motor impairments, including tremors, muscle rigidity, and slowness of movement. However, the progression of the disease is also associated with the development of nonmotor symptoms, including cognitive deficits and affective disorders (Aarsland et al., 2010; Martinez-Martin et al., 2007; Riedel et al., 2010). The cognitive symptoms of PD are attributed to impaired function of the frontal cortex (Mihaescu et al., 2019; O’Callaghan et al., 2014; Sawada et al., 2012), possibly resulting from depletion of monoamine neurotransmitters and acetylcholine (Buddhala et al., 2015; Halliday et al., 2014; Mattila et al., 2001; Riekkinen et al., 1998), as well as neurodegeneration affecting the cortex in the final stages of the disease (Armstrong, 2017; Braak et al., 2003; González-Redondo et al., 2014). The impaired function of the frontal cortex also contributes to the motor symptoms of PD due to altered activity and the loss of specificity of efferent neurons (Rowe et al., 2002; Vercruysse et al., 2014). The cellular mechanisms associated with impaired frontal cortex activity remain only partly understood. A number of studies have assessed changes in the transcriptome and proteome in prefrontal cortex samples derived *postmortem* from PD patients and in some cases supplement the results with large-scale analysis of single nucleotide polymorphisms (Duke et al., 2006; Dumitriu et al., 2016, 2012; Hoss et al., 2016; Riley et al., 2014). Extensive changes in either transcription or protein abundance were observed, including differences in the expression of not only genes encoding metallothioneins, the transcription factor *FOXO1* and its network of regulated genes but also mitochondrial genes, proteasome components, and multiple short RNAs. Nevertheless, the reported results are not in clear consensus with regard to the underlying mechanisms. Additionally, most of the analyses were based on samples from patients with late stages of the disease who had received extensive treatment. Thus, it is difficult to distinguish whether the observed effects should be attributed to late-stage neurodegeneration, neurotransmitter depletion or the effects of long-term medication.

The primary treatment of PD relies on L-3,4-dihydroxyphenylalanine (L-DOPA), a dopamine precursor. L-DOPA is effective in alleviating motor impairments, especially in the initial period of treatment, but displays mixed efficacy against the nonmotor symptoms of PD. For example, L-DOPA was observed to improve working memory (Simioni et al., 2017) and cognitive flexibility but simultaneously increased impulsivity (Cools et al., 2003) and failed to restore impaired sequence learning (Ghilardi et al., 2007) in PD patients. Moreover, the administration of L-DOPA altered frontal cortex connectivity in healthy volunteers and had psychoactive effects, increasing impulsivity and altering performance in tasks dependent on executive functions (Kelly et al., 2009; Shiner et al., 2015). The molecular adaptations involved in the effects of L-DOPA on the frontal cortex remain elusive.

A model commonly used to study the effects of dopamine depletion on brain physiology is the 6- hydroxydopamine (6-OHDA)-induced lesion of dopaminergic neurons (Simola et al., 2007; Ungerstedt, 1968). Lesioning is usually performed unilaterally, which permits comparison of the functioning of a dopamine-depleted hemisphere and an intact hemisphere. 6-OHDA lesioning is widely used as it reproduces a PD-like phenotype resulting from damage to the ventral midbrain neurons and involves the disruption of mitochondrial activities that mediate cytotoxic processes (Bezard et al., 2013; Kreiner, 2015; Simola et al., 2007); thus, 6-OHDA lesioning has potential construct validity as a model of human PD (Hauser and Hastings, 2013). Several studies have examined the effects of 6-OHDA lesions, in some cases followed by L-DOPA treatment, on gene expression in the forebrain and striatum in particular (Berke et al., 1998; Heiman et al., 2014; Konradi et al., 2004; Smith et al., 2016). These studies revealed that neurons on the lesioned side of the striatum showed changes in the expression of immediate early-response genes (IEGs), *Grin1* (an NMDA receptor subunit), *Stx6* (involved in exocytosis) or *Ldhb* (metabolism). To the best of our knowledge, no studies have comprehensively examined transcriptomic changes evoked by L-DOPA in the frontal cortex.

Here, we investigated the effects of L-DOPA on gene expression in the frontal cortex of rats with unilateral 6-OHDA lesions of the ventral midbrain neurons. We analyze L-DOPA-regulated gene expression in the contexts of potential regulatory mechanisms, cell-type specificity, and similarities to the transcriptional signatures of other drugs.

## Materials and methods

### Animals

The tissue samples used in this study were derived from a previously reported experiment (Lorenc-Koci et al., 2013). All the previously described animal procedures were performed in accordance with the European Union guidelines for the care and use of laboratory animals (Directive 2010/63/EU) and had been approved by the Second Local Institutional Animal Care and Use Committee in Krakow (permit No. 846/2011). Briefly, adult male Wistar Han rats (Charles River, Germany) with an initial body weight of 300 g ± 20 g were kept in the Maj Institute of Pharmacology animal facility (5 animals per cage) under standard laboratory conditions (22 °C, light/dark cycle of 12/12 hours with lights on beginning at 7 a.m.). Animals had unlimited access to food and water. The rats were unilaterally lesioned by the infusion of 6-OHDA hydrochloride (8 µg, free base) into the left medial forebrain bundle. The animals received desipramine hydrochloride (i.p., 25 mg/kg) prior to surgery to prevent damage to noradrenergic neurons. After the surgery, the rats were allowed to recuperate for 2 weeks, and then the extent of the lesion was verified using the apomorphine-induced rotation test. The inclusion criterion for further experiments was ≥ 98 contralateral turns in one hour. On the day following the rotation test, rats started a 14-day treatment period with L-DOPA (12.5 mg/kg, i.p., once daily) combined with benserazide hydrochloride (6.25 mg/kg, i.p., 30 minutes prior to L-DOPA). The control animals received a 0.9% (w/v) NaCl solution. Rats were killed by decapitation 1 hour after receiving the last dose of L-DOPA or saline.

### RNA sequencing

Frontal cortex samples from both hemispheres were dissected separately on an ice-chilled plate, using the olfactory bulbs and the most anterior bifurcation of the middle cerebral artery as orientation points to make cuts in the frontal plane. Striatal tissue was removed from the sections, and the cortices were separated into the left and right sides and stored at -80°C before proceeding. Tissue samples were homogenized using a TissueLyser (QIAGEN, Germany), and total RNA was extracted with an RNeasy Mini Kit (QIAGEN) using a QIAcube (QIAGEN) in accordance with the manufacturer’s protocol. The integrity and concentration of the extracted RNA were assessed using an Agilent RNA 6000 Nano Kit on a 2100 Bioanalyzer (Agilent, USA). Based on RNA integrity number (RIN > 7.2) values, samples from 5 animals per group were chosen for RNA sequencing. RNA sequencing was performed as an external service by Novogene (Hong Kong). Briefly, poly(A) RNA was isolated from the total RNA samples by the addition of oligo(dT) beads. cDNA libraries were prepared on a template of randomly fragmented mRNA using the NEBNext Ultra II Directional RNA Library Prep Kit for Illumina (New England Biolabs, USA). Since 8 samples out of 20 initially contained < 200 ng of total RNA, all the samples were processed to prepare cDNA from poly(A) RNA following the low-input protocol. Complete cDNA libraries consisting of reads 250-300 base pairs in length acquired from the samples were subjected to sequencing on an Illumina system (150 base pairs, paired-end, 20 million reads per sample).

### Data preprocessing and differential expression analysis

A quality check of the raw RNA-seq data was performed with FastQC v0.11.8 in R 3.4. The reads were aligned to the Rnor6.0 rat reference genome from the Ensembl database using HISAT2 v2.1.0 (Kim et al., 2019). Transcript counts were normalized to fragments per kilobase of transcript per million fragments mapped (FPKM) values with the Cufflinks v2.2.1 package (Trapnell et al., 2010). We used two-way ANOVA with lesion as a within-subject factor and treatment as a between-subject factor on log2(1 + FPKM) values for each gene to detect statistically significant differences. The Benjamini-Hochberg false discovery rate (FDR) correction for multiple comparisons was used to adjust *P*-values. Statistical significance testing was performed on transcripts with mean log2(1 + FPKM) values greater than 1. The BioMart interface to the Ensembl database was used for annotation of the transcripts. Hierarchical clustering was performed using distances calculated as (1- Pearson’s R^2^). Genes with FDR values smaller than 0.05 for the L-DOPA treatment effect are referred to as “differentially expressed genes”.

### Identification of coexpression networks

Weighted gene coexpression network analysis was performed with the WGCNA package v1.68 (Langfelder and Horvath, 2008) in R v3.6.1, and was applied with a step-by-step approach to construct coexpression networks from genes with mean log2(1 + FPKM) values greater than 1. Briefly, a soft-threshold power of 11 was set based on scale-free topology calculations. Then, adjacency values were transformed into a topology overlap matrix (TOM) with TOMtype = “signed”, and TOM-based dissimilarity (dissTOM) was defined as 1-TOM. A network dendrogram of genes was constructed based on average linkage hierarchical clustering and dissTOM with the “Dynamic Tree Cut” method and the minimum module size set to 30. The resulting gene modules were assigned colors for easier reference, with the “gray” module comprised of genes that did not match the inclusion criteria for any other module. Module eigengene values were clustered based on their correlation, and modules identified by cutreeDynamic were merged based on the threshold MEDissThres = 0.5 by the mergeCloseModules function.

### Transcription factor binding

The Seqinspector tool was used to assess overrepresentation of transcription factor-binding sites in the promoter regions of differentially expressed genes (Piechota et al., 2016). Seqinspector uses datasets from ChIP-seq experiments performed in mouse or human cells; thus, we translated the list of rat genes to their mouse homologs and used the mm10 mouse genome assembly as background. Rat genes in the lists were translated to mouse homologs with biomaRt v2.40.5 (Durinck et al., 2005). The following rat genes were excluded due to the existence of more than 1 or no mouse homologs or differences in the symbols’ names: *Il6r, Nfkbia* in the case of the differentially expressed genes, and *Nfkbia, Nat8f3, Gpr52, RGD1311899, Rcor2l1, Spag5, LOC100912481*, and *LOC100911313* in the case of the “salmon” module genes. The resulting records from corresponding sequencing experiments (tracks) were considered statistically significant if their Bonferroni-corrected *P*-values were smaller than 0.05.

### Cell-type specificity of gene expression

Analysis of the brain cell subtypes in which transcripts of the differentially expressed genes were present was performed using “RNA-Seq Data Navigator” from the Cell Types Database (Allen Institute for Brain Science 2015). The database is based on RNA-seq profiling of mouse cortical cells, and we used the list of murine homologs of the rat genes indicated by ANOVA as described above. The *Dipk2a, Noct* and *Cavin2* mouse genes were not recognized by the database and excluded from the analysis, along with *Nfkbia* and *Il6r*. The result of this analysis was the prevalence of transcripts of our differentially expressed genes within different cell subtypes (Tasic et al., 2018). The exported group fraction values for each gene were then used to plot a heatmap.

### Annotation enrichment analysis

The differentially expressed and “salmon” cluster genes were analyzed for annotation enrichment using the enrichR package v2.1, an R interface to the Enrichr web server (Chen et al., 2013; Kuleshov et al., 2016). Enrichr uses the human genome as the reference set; thus, we used biomaRt v2.40.5 to find human homologs of the rat genes. Genes with different symbols, genes without a human homolog, and genes that had more than 1 human homolog were excluded from the analysis. The following genes were excluded: *Zfp189* and *Nfkbia* from the list of differentially expressed genes and *RGD1311899, Zfp189, Nat8f3, Nfkbia, Gpr52, Sik1, Zfp521, Rcor2l1, Spag5, LOC100912481, LOC100911313* from the list of genes included in the “salmon” WGCNA module. The results were considered significant at an adjusted *P-*value smaller than 0.05.

## Results

### L-DOPA-induced gene expression

Experiments were performed on frontal cortex samples derived from a previously described group of animals that underwent unilateral lesion of dopaminergic neurons with 6-OHDA followed by 14 days of L-DOPA treatment (Lorenc-Koci et al., 2013). A diagram summarizing the procedure is shown in Figure 1. We dissected the left and right frontal cortices and separately isolated total RNA from each of them. For both treatment groups, treated with saline or L-DOPA, we analyzed 5 paired samples (left and right cortices from the same rat, 20 samples in total). RNA sequencing of poly(A)-enriched cDNA yielded on average 26.4 million pairs of raw reads per sample. The sequence reads were mapped to the Ensembl rat genome assembly Rnor6.0 using HISAT2. The reads were aligned to a total number of 33,883 sequences, and counts were normalized to FPKM values. There were 12,455 genes with a mean log2(1 + FPKM) value greater than 1. These genes included 12,034 protein-coding genes, 259 pseudogenes, 98 long intergenic noncoding RNAs, 2 mitochondrial rRNAs, 1 nucleolar rRNA, 2 ribozymes, 3 scaRNAs, and 1 snoRNA (for complete results, see Table S1).

**Figure 1.**
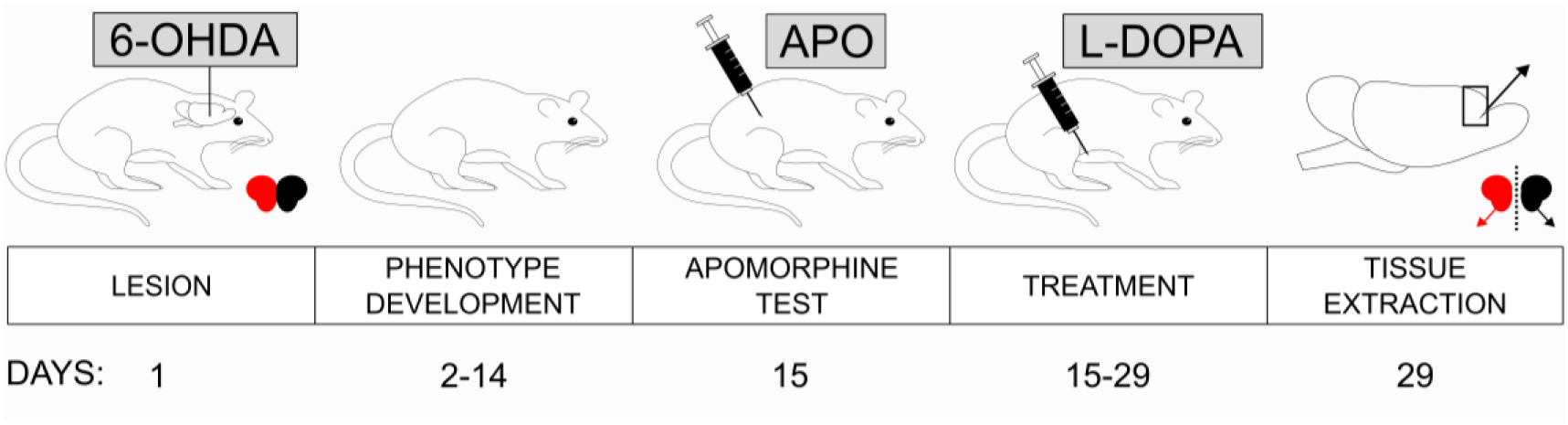
Treatment scheme of rats with unilateral lesions of dopaminergic neurons as a model of Parkinson’s disease. Male adult rats were infused with 8 μg/4 μl 6-hydroxydopamine (6-OHDA) into their left medial forebrain bundle. The toxin was used to kill dopaminergic neurons to produce an animal model of Parkinson’s disease. Two weeks later, the animals were injected s.c. with 0.25 mg/kg apomorphine (APO) to induce rotational behavior. The number of rotations in 1 hour was scored, and only the animals that exhibited at least 98 contraversive rotations (as an indication of sufficient nigral degeneration) were used in the following procedures. One day after the APO test, the treatment regime started. Control rats received i.p. saline, while the experimental group received i.p. benserazide (6.25 mg/kg) once daily for 14 days, followed by L-DOPA (12.5 mg/kg). The animals were decapitated 1 hour after the last dose of saline or L-DOPA, and their frontal cortex tissue was individually dissected from each hemisphere (the procedure was previously described and published by Lorenc-Koci et al. (2013)).

To identify differentially expressed genes, we performed two-way ANOVA with FDR correction for multiple comparisons. Out of 12,455 genes, there were 48 genes with FDR values smaller than 0.05 for the “treatment” factor, as summarized in Figure 2 and Table S2. No genes were found to be significantly differentially expressed between the lesioned (left) and nonlesioned (right) frontal cortices, and no statistically significant interactions between effects were observed (Table S2). L-DOPA treatment caused an increase in the abundance of transcripts corresponding to 38 genes (upper part of the heatmap) and downregulated 10 genes (lower part). All the differentially expressed genes were protein-coding genes and had functions related to the immune response (*Bcl6, Ifngr1, Il6r*), modification of the extracellular matrix (*Hyal2, Mmp9*), neuronal signal transduction (*Chrm4*), the circadian rhythm (*Per1*), cellular uptake (*Tfrc, Slc2a1*), the stress response (*Sgk1*), cell differentiation (*Sox2, Nedd9*), and the response to hypoxia (*Ddit4*), among others. Moreover, we noted that several genes, including *Per1, Sgk1, Errfi1, Id1*, and *Klf4*, were associated with the immediate early response. Mean transcript abundancies of the differentially expressed genes normalized as log2(1 + FPKM) were in the range of 1.16 (*Mmp9*) to 6.75 (*Tspan17*).

**Figure 2.**
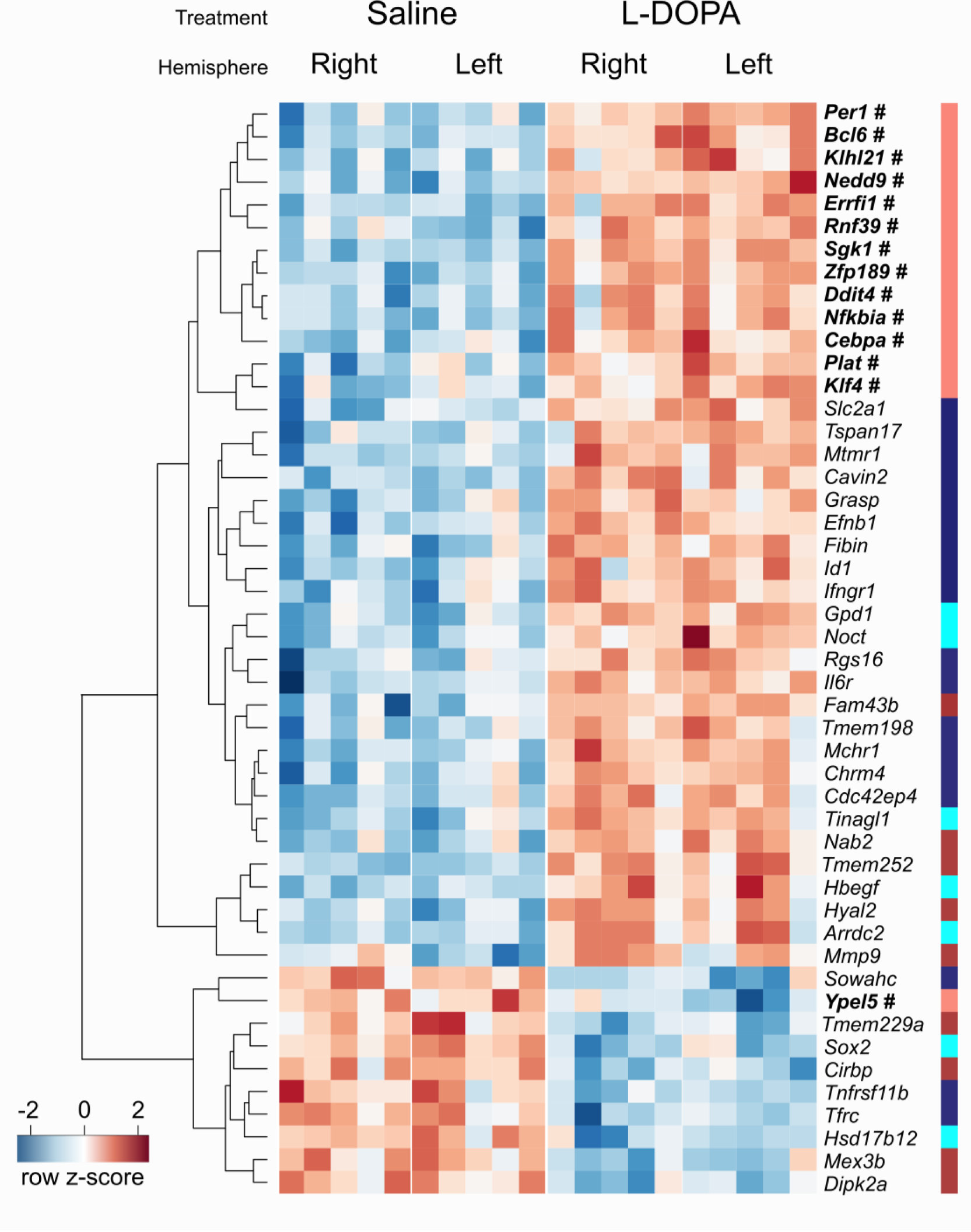
Gene expression changes evoked by L-DOPA in the frontal cortex of rats with unilateral lesions of the dopaminergic system. A total of 48 genes were found to have altered transcript abundance after L-DOPA treatment in the frontal cortex of rats with unilateral 6-hydroxydopamine lesions (two-way ANOVA, FDR < 0.05). Each column represents one sample (left or right frontal cortex), and rows correspond to genes, as indicated on the right. Colors represent normalized expression levels. Genes are ordered based on hierarchical clustering, and the dendrogram is shown on the left. Additionally, the colored stripe on the right shows the assignment of genes to modules from the WGCNA. Genes belonging to the “salmon” module are highlighted in bold and marked with a “#”.

### Identification of coexpression networks

To identify networks of coexpressed genes, we employed weighted gene coexpression network analysis (WGCNA). This method addresses some of the weaknesses associated with the identification of differentially expressed genes by null hypothesis testing, such as arbitrary significance criteria or the assumption that the expression of individual genes is independent. We performed step-by-step WGCNA with a soft-threshold power of 11, the lowest number for which the scale-free topology R^2^ was greater than 0.8. The 12,455 genes were clustered with the “Dynamic Tree Cut” method, which resulted in the detection of 17 modules, each labeled with a color (Figure 3A). There were 5 modules detected at the merging threshold of 0.5 (Table S3): “brown”–7148 genes (including 6909 protein-coding genes), “cyan”–1450 genes (1401), “gray”–287 genes (257), “midnight blue”–3490 genes (3388) and “salmon”–80 genes (79; the assignment of genes to modules is listed in Table S1). The modules in the figure are not continuous, as it was not possible to accurately represent distances based on eigenvalues in a two-dimensional plot.

**Figure 3.**
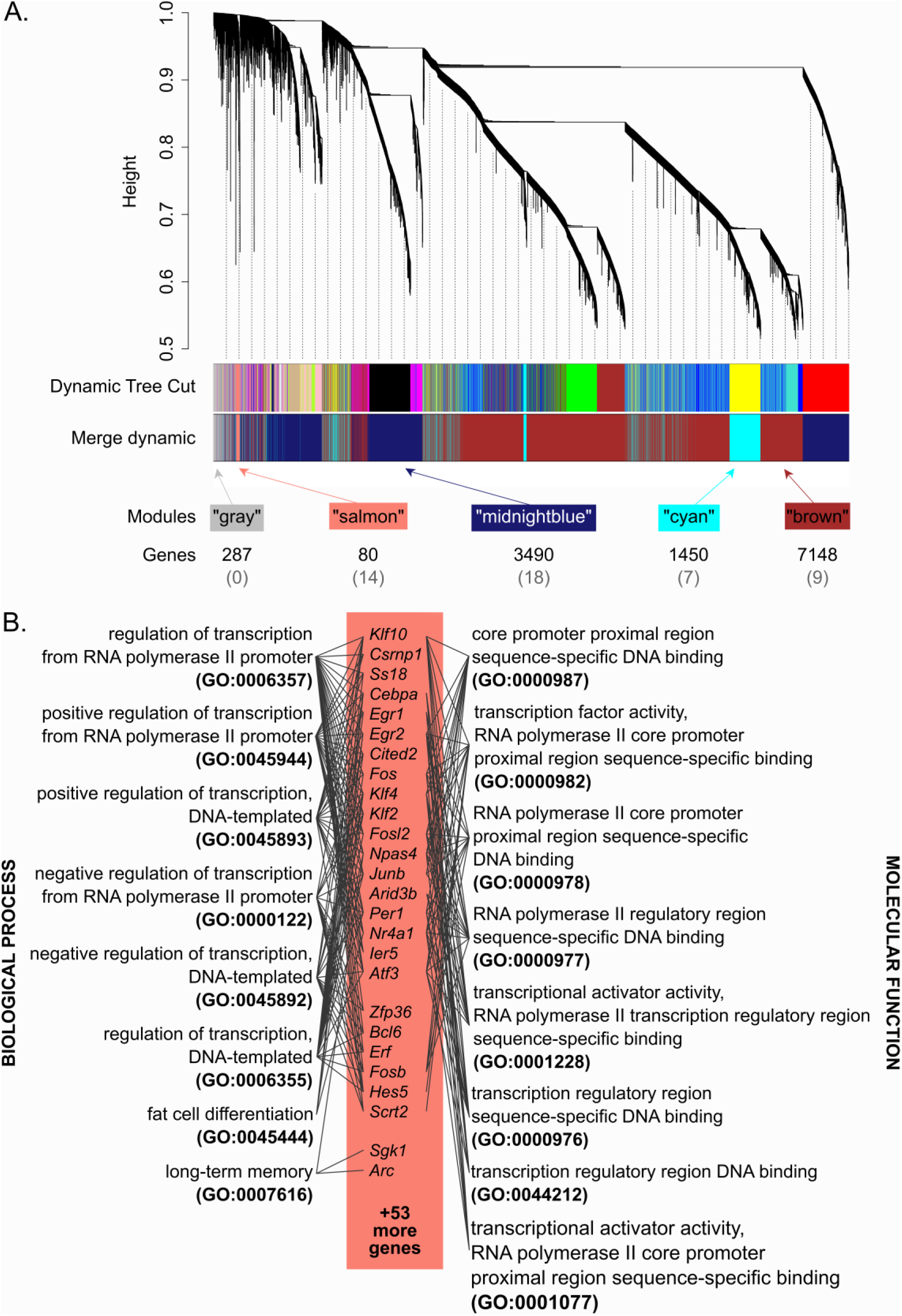
L-DOPA-regulated gene coexpression networks in the rat frontal cortex. (A) Graphic representation of gene assignment of the 12,455 genes (with log2(FPKM + 1) values greater than 1) to weighted gene coexpression network analysis modules. Each branch of the dendrogram and colored stripe represent one gene, and the colors indicate assignment to a particular module named by color. The ‘Dynamic Tree Cut’ method detected 17 modules (the upper block of colored stripes), which were further joined with the ‘Merge dynamic’ method at a threshold of 0.5 into 5 resulting modules (the lower block). The numbers below the blocks indicate the total number of genes in a module, and those in brackets show the number of genes in the module that overlap with the differentially expressed genes. (B) The “salmon” module included many immediate early genes and was enriched in gene ontology (GO) terms related to the process of transcription. Biological process GO terms are on the left, and molecular function terms are on the right. Gray lines connect the terms with the corresponding genes.

The smallest of the identified clusters, the “salmon” cluster, had high overlap with the results of ANOVA and included 14 out of the 48 differentially expressed genes (Figures 2 and 3A, Table S2). Seventy-nine out of 80 genes in this module were protein-coding genes (one was a pseudogene), and the module included several IEGs (e.g., *Fos, Fosb, Junb, Egr1, Klf4, Atf3, Arc*, and *Npas4*), a large proportion of which were related to transcription processes (Figure 3B). Of the 14 overlapping genes, 13 were clustered in the dendrogram in Figure 2. The proximity of these 13 genes on the heatmap is in line with expectations, as clustering is based on an approach similar to that of the identification of coexpressed modules. The overlap in results from the two analytical methods crossvalidate the effects on L-DOPA on transcripts associated with immediate early gene expression, which was hemisphere-independent (no specific effect of the lesion on the ipsilateral side).. The other coexpression modules had relatively low overlap with the differentially expressed genes (Figure 3A). The “midnightblue” module contained 18 differentially expressed genes: 15 of which were upregulated and 3 of which were downregulated. Of these 15 genes, 9 genes (*Slc2a1, Tspan17, Mtmr1, Cavin2, Grasp, Efnb1, Fibin, Id1* and *Ifngr1*) were adjacent in the heatmap (Figure 2). Additionally, the remaining 6 upregulated genes were clustered on the dendrogram (branches containing *Tmem198, Mchr1, Chrm2* and *Cdc42ep4*, and *Rgs16* and *Il6r* individually). We noted, however, that an overlap of 18 among 3490 is low and probably negligible. Even smaller overlaps were observed in the case of the “brown” (9 genes among 7148) and “cyan” (7 genes among 1450) modules. The “gray” module, a collection of genes that were not allocated to any module of coexpressed genes, did not include any differentially expressed genes. Therefore, we performed further analyses of gene promoters and annotation enrichment on the “salmon” module to obtain a result complementary to the set of differentially expressed genes.

### Cell-type specificity of gene expression

We used large-scale single-cell RNA-seq datasets from the Allen Institute for Brain Science to identify cell types with previously confirmed transcription of the differentially expressed genes (Allen Institute for Brain Science, 2015; Tasic et al., 2018). For each gene of interest, we queried the fraction of cells of each type in which it had been found to be present (the criterion in the original paper was CPM ≥ 1.). The reported cell classification was based on the expression of marker genes and clustering of transcriptomic profiles using WGCNA and principal component analysis. The results of the analysis are summarized as a heatmap shown in Figure 4. The use of fractions permits the assessment of the ubiquity of gene expression, although it is not necessarily indicative of high levels of transcription. In general, the heatmap shows that the differentially expressed transcripts vary greatly in the ubiquity of their expression. The upper part clusters transcripts present in most types of cells and essentially all types of neurons. *Ypel5* is an example of a differentially regulated transcript that was highly ubiquitous (fraction of cells > 0.33 across all the cell subtypes). A group of differentially expressed transcripts was prevalent in glutamatergic and GABAergic neurons, i.e., *Errfi1, Fam43b, Mtmr1, Tmem198* and *Tspan17*. Genes with relatively specific expression restricted to particular subpopulations of GABAergic neurons (i.e., parvalbumin or serpin F positive) were *Hbegf* and *Rgs16*, while *Rnf39* expression was detected mainly in glutamatergic neurons. None of the differentially expressed genes were specifically expressed by a single type of neuron. The most specific expression was observed in the cases of *Cebpa* and *Klf4*, which appeared restricted to macrophages and endothelial cells, respectively. Some transcripts, such as *Mmp9* and *Tnfrsf11b*, were detected in fewer than one-third of cells in all subtypes. Taken together, these results show that the observed changes in expression are not restricted to a singular type or subtype of cells and may not necessarily be primarily related to neurons.

**Figure 4.**
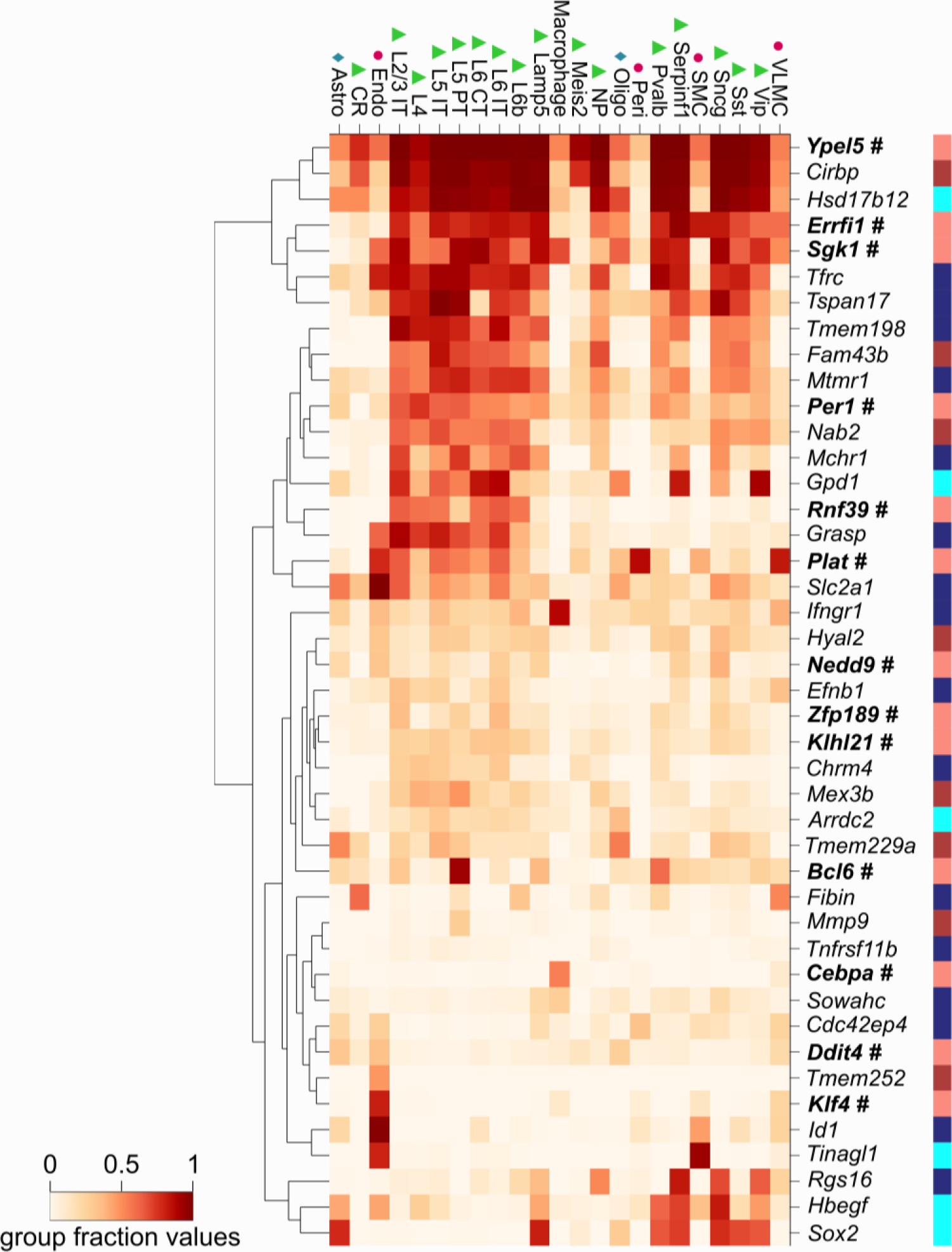
Analysis of cell-type specificity of the expression of genes differentially expressed in the rat frontal cortex after L-DOPA treatment. The heatmap includes genes that passed the differential expression criterion and had mouse orthologs. Each column reflects a cell subtype, based on the Allen Atlas classification, and rows correspond to genes. The color of the squares in the heatmap indicates the fraction of the cells belonging to the subtype that were found to express a particular gene (legend shown in the bottom left). Symbols above the heatmap mark broader cell categories: triangles– neurons, diamonds–glia, and dots–cells associated with the vascular system. The colored stripe on the right shows the assignment of genes to modules from the WGCNA. Genes belonging to the “salmon” module are highlighted in bold and marked with a “#”. Abbreviations: Astro – astrocytes; CR – Cajal- Retzius cells; CT – corticothalamic neurons; Endo – endothelial cells; IT – intratelencephalic neurons; L – cortical layer; NP – near-projecting neurons; Oligo – oligodendrocytes; Peri – pericytes; PT – pyramidal tract neurons; SMC – smooth muscle cells; VLMC – vascular leptomeningeal cells.

### Gene promoter and annotation analyses

We analyzed the promoter regions of genes with differential expression and the “salmon” module for transcriptional factor binding using Seqinspector. The analysis identified 8 transcription factors with significantly increased binding in the promoter regions of the differentially regulated genes: early growth response protein 1 (EGR1), EGR2, glucocorticoid receptor (GR), hypoxia-inducible factor 1- alpha (HIF1A), myoblast determination protein 1 (MYOD), myogenin (MYOG), neuronal PAS domain protein 4 (NPAS4), and transcription factor 3 (TCF3; Table S4). In the case of the “salmon” module genes, the analysis identified 7 transcription factors, namely MYOG, GR, NPAS4, and also serum response factor (SRF), CCAAT/enhancer-binding proteins beta and delta (CEBPB and CEBPD, respectively), CREB-binding protein (CBP; the complete list is presented in Table S5.). The results are consistent with previous analyses showing that EGR1, EGR2, NPAS4 and SRF are involved in activation of the expression of IEGs (Benito and Barco, 2015; Herdegen and Leah, 1998). Additionally, we previously observed that GR-dependent expression appears in the acute response to treatment with most drugs, including psychostimulants and drugs acting on dopamine receptors (Piechota et al., 2012; Zygmunt et al., 2019). The results of ontology analysis of the genes assigned to the “salmon” module with enrichR also appear consistent with the idea of the activation of IEGs. A large proportion of these genes act as transcription regulators, and accordingly, 6 out of 8 enriched GO biological process terms, as well as 8 out of 8 enriched GO molecular function terms, were related to transcriptional processes (Figure 3B; for full results, see Table S6). One of the GO biological process terms with significant overrepresentation was “long-term memory” (GO:0007616), and the term was assigned to *Npas4, Arc* and *Sgk1*. On the other hand, the 46 differentially expressed genes tested were not significantly enriched in GO terms.

Next, we assessed overlaps between L-DOPA-regulated genes and gene sets related to the response to pharmacological treatment in the following databases: the Drug Signatures Database (Yoo et al., 2015) and the Gene Expression Omnibus (GEO) drug perturbations database (full analysis results are shown in Table S7.). A large number of significant overlaps were identified, including expression patterns associated with steroids (e.g., dexamethasone, estradiol, corticosterone), antidiabetics (rosiglitazone, insulin), cytotoxic and cytostatic agents (e.g., vemurafenib, actinomycin D), vitamins (e.g., K3, retinoic acid, ascorbic acid), drugs targeting immune responses (e.g., lipopolysaccharide, etanercept), toxins (e.g., sodium arsenite, vanadium pentoxide, benzene), and drugs of abuse (morphine, heroin, cocaine, ethanol). The results of analysis of the “salmon” module genes were largely similar (Table S8), with additional significant results from the DrugMatrix Database (National Toxicology Program, 2010) that did not reveal any significant overlap with the list of differentially expressed genes. Importantly, these transcriptional signatures were reported in different organisms and tissues, including the rat liver, a human breast adenocarcinoma cell line and the murine striatum. We noted that 41 of the L-DOPA-regulated genes and 65 of the genes in the “salmon” module were represented in both drug-related expression signature databases (i.e., Drug Signatures Database and GEO database). This indicates that a large subset of genes influenced by L-DOPA are also activated by multiple other drugs, which may indicate a nonspecific response. For instance, among the 319 significant overlaps between the sets from all the drug signature databases and the 46 differentially expressed genes used in the analysis, *Ddit4* appeared 155 times, and *Hbegf* appeared in 114 cases (Table S7). The same was the case for genes in the “salmon” module: *Atf3* appeared in 724 out of 1166 total drug-regulated transcription signatures, 594 signatures included *Egr1*, and 421 signatures listed *Dusp1* (Table S8). Conversely, 34 of the differentially expressed genes were present in the enrichment results of all the relevant drug-related databases, and among genes in the “salmon” module, 19 were present in the sets from the “salmon” significantly overrepresented databases. The 34 differentially expressed genes in the drug sets were not associated with any of the specific cell subtypes, based on the analysis whose results are shown in Figure 4. These results may suggest that genes not included in the overlaps with the transcriptional signatures of other drugs are associated with molecular mechanisms specific to L-DOPA action. The 4 genes that were not present in the transcriptional signatures were *Grasp, Dipk2a, Noct*, and *Cavin2*. However, the latter three were not recognized by the Allen Institute for Brain Science’s Cell Types Database analytical tool. This leaves *Grasp*, which appears to be expressed predominantly in neurons as well as endothelial cells (Figure 4).

Finally, we performed a direct comparison of the results of gene expression analysis with those of a recent comprehensive analysis of L-DOPA-induced transcription in the rat striatum (Smith et al., 2016). A previous study identified 95 genes with decreased abundance in striatal tissue in a unilateral 6-OHDA rat model of L-DOPA-induced dyskinesia, of which 19 were included in the “salmon” module (e.g., *Arc, Atf3, Egr1, Egr2*, and *Nr4a1*; for a full list, refer to Table S9.) and 4 in the list of the differentially expressed genes (i.e., *Klf4, Per1, Cdc24ep4*, and *Nab2*). However, it should be noted that there are important methodological differences between the cited study and this report, including the use of a different dose of L-DOPA (4 mg/kg) in the case of the experiments reported by Smith and colleagues (2016).

## Discussion

We found that L-DOPA treatment in rats with unilateral lesions of dopaminergic neurons induced bilateral changes in gene expression in the frontal cortex. The differentially expressed transcripts are functionally diverse and include genes engaged in the immediate early response of the cell. Analysis of the cell-type specificity of gene expression indicated that transcription changes occurred in both neuronal and nonneuronal cell types. Finally, we found that the differentially expressed genes, with few exceptions, overlapped with genes regulated by other, functionally unrelated drugs.

6-OHDA-induced lesioning of dopamine neurons in rats is a widely used model of PD (Simola et al., 2007). Several studies demonstrated differences in IEG induction following administration of a D1- like receptor agonist and the contents of dopamine and its metabolites between the lesioned and nonlesioned hemispheres in animals with unilateral 6-OHDA lesions, particularly in the striatum (Berke et al., 1998; Perese et al., 1989; Walker et al., 2013), and in the level of monoamine neurotransmitters in the substantia nigra, hippocampus and frontal cortex (Kamińska et al., 2017). Unexpectedly, in this study, we found no significant differences in gene expression between the lesioned and intact sides of the frontal cortex. Thus, either the unilateral lesion had no appreciable basal effect on homeostasis in the frontal cortex, or the effects on both sides were the same. We would argue that the former is more likely for the following reasons. First, our previous results confirmed depletion of dopamine on the lesioned side of the frontal cortex of rats that underwent the same treatment (Kamińska et al., 2017). Second, the expression of markers of glial activation (e.g., *Gfap* and *Aif1*) appeared normal and not indicative of glial proliferation. This would imply a major difference from the observed sensitization of dopamine receptor D1 signaling observed in the dopamine-depleted striatum (Berke et al., 1998). Moreover, a comparison of the effects of L-DOPA on gene expression with results reported in the case of the striatum (Smith et al., 2016) shows some overlap in the differentially regulated genes identified. However, the changes in transcription appear opposite in the case of several putative IEGs. This could be consistent with the notion that L-DOPA treatment in the dopamine-depleted forebrain increases excitatory transmission from the thalamus to cortical areas but reduces excitatory inputs from the cortex to the striatum (De Deurwaerdère et al., 2017). Furthermore, we note that the previous proteomic and transcriptomic analyses performed on samples taken *postmortem* from PD patients did indicate neurodegeneration-related gene expression in the prefrontal cortex (Dumitriu et al., 2016). Conversely, there was no indication of frontal cortex degeneration after 6-OHDA lesioning in rats (Simola et al., 2007), which may in part explain the lack of differences in gene expression between hemispheres that we observed. Finally, 6-OHDA lesioning causes the loss of dopaminergic neurons within days of treatment, while in PD patients, degeneration develops over years before the onset of motor impairments (Morrish et al., 1998). Thus, while there is overlap to some extent in gene expression changes reported here and in previous studies on *postmortem* cortex samples, e.g., *Zfp189, Klhl21, Bcl6*, and *Dddit4* (Dumitriu et al., 2016) or *Nab2* and *Sox2* (Dumitriu et al., 2012), a direct comparison of 6-OHDA lesioning with *postmortem* analyses does not appear valid. We consider this to be the primary limitation of our study.

Our data showed that L-DOPA treatment had a robust effect on gene expression in the frontal cortex. The effects of L-DOPA appeared to involve two components: a general response and a drug-specific response. Both would be of relevance in the context of the mechanisms through which L-DOPA treatment affects cognitive functions requiring cortical activity. We note extensive similarities between changes in gene expression induced by L-DOPA treatment and those induced by other drugs, but we cannot exclude the possibility that these changes are specific to select cell types. Thus, the specific aspect of drug-induced gene expression could be the profile of the cells affected, rather than the exact list of affected transcripts. Independently, as shown in the results, a single induced transcript, *Grasp* (General Receptor For Phosphoinositides 1 Associated Scaffold Protein, also known as Tamalin), appeared to be specific to the effects of L-DOPA. Caution should be taken in the case of such singular results, but we note that the Grasp protein was reported to be involved in the trafficking of metabotropic glutamate receptors (Kitano et al., 2002), and its increased abundance was reported in *postmortem* CA1 hippocampal region samples from persons diagnosed with schizophrenia compared to healthy controls (Matosin et al., 2014). *Grasp* has also been shown to be upregulated in the ipsilateral striatum of rats with 6-OHDA unilateral lesions following chronic L-DOPA exposure, as well as in Flinders-resistant line rats that developed abnormal involuntary movements (a symptom of L-DOPA-induced dyskinesia) (Schintu et al., 2020). These observations support the notion that *Grasp* could be part of a mechanism involved in the effects of L-DOPA on cognitive functions, though this remains a conjecture.

An additional point to consider is the cellular selectivity of the response to L-DOPA. The method employed here to determine the cell-type specificity of gene expression does not provide direct information on cell-level gene expression changes; rather, we extrapolate data based on the fraction of neuronal and nonneuronal cells that were observed to express the genes of interest. The analysis suggests a heterogeneous response to L-DOPA in the frontal cortex involving different subtypes of neurons: glia, macrophages and vascular cells. The caveat is that these changes are likely occurring in only subsets of the cells indicated and speculatively may even involve lesion- or L-DOPA-induced transcription in cells that normally do not express the differentially regulated genes.

Taken together, the results of gene expression analysis in the context of previously reported data lead to two conclusions. First, the effects of L-DOPA on the frontal cortex are not lateralized and thus differ from reported transcription changes in the striatum. Second, the genes affected by L-DOPA overlap with those reported in the cases of several other drugs; hence, the key feature associated with the clinical effects of L-DOPA is probably cell-type specificity. We believe that extending the analysis of the effects of L-DOPA on gene expression to achieve both cell type and anatomic resolution could lead to the identification of patterns associated with antiparkinsonian efficacy.

## Data availability

RNA-seq project data: NCBI BioProject (accession No. PRJNA547879) www.ncbi.nlm.nih.gov/bioproject/?term=PRJNA547879

RNA-seq raw data: NCBI Sequence Read Archive (accession No. SRX6071775 - SRX6071794) www.ncbi.nlm.nih.gov/sra?linkname=bioproject_sra_all&from_uid=547879

Scripts:

github.com/ippas/ifpan-annaradli-ldopa

## Abbreviations

6-OHDA: 6-hydroxydopamine;
APO: apomorphine;
IEGs: immediate early genes;
L-DOPA: L-3,4-dihydroxyphenylalanine;
PD: Parkinson’s disease;
WGCNA: weighted gene co-expression network analysis

## Acknowledgements

The authors declare no conflict of interest.

## Authors’ contribution

In accordance with the CRediT roles, the authors contributed to this work as follows: AR – Investigation, Formal analysis, Data curation, Writing – Original draft preparation, Visualization; KK – Investigation, Resources, Writing - Review & Editing; MB – Formal analysis, Data curation, Writing - Review & Editing; MP – Formal analysis, Data curation; MK – Formal analysis, Data curation, Writing - Review & Editing; JP – Supervision, Writing - Review & Editing; ELK – Conceptualization, Investigation, Supervision, Resources, Funding acquisition; JRP – Conceptualization, Investigation, Formal analysis, Supervision, Writing – Original draft preparation, Visualization.

## Supplementary materials’ legends

**Table S1**. Sequencing results. Genes mapped to the rat Rnor_6.0 genome assembly, with Ensembl IDs, gene symbols, mean and individual log2(FPKM +1) values, false discovery rate (FDR) values for both factors and their interaction, gene classification, and the WGCNA modules to which the genes were assigned.

**Table S2**. List of differentially expressed genes, with Ensembl IDs, gene symbols, mean log2(FPKM +1) values, false discovery rate (FDR) values for all factors, fold changes for each contrast, and the WGCNA modules to which the genes were assigned.

**Table S3**. Module eigengenes (MEs) of the five WGCNA modules after merging the modules identified by the “Dynamic Tree Cut” method. The columns indicate modules, and the rows represent samples.

**Table S4**. Enrichment of transcription factor-binding sites in the loci of the differentially expressed genes.

**Table S5**. Enrichment of transcription factor-binding sites in the loci of genes assigned to the “salmon” WGCNA module.

**Table S6**. Significantly enriched gene ontology (GO) terms from the biological process and molecular function branches in the list of genes assigned to the “salmon” WGCNA module.

**Table S7**. Significant overlaps between differentially regulated genes and transcriptomic signatures from Gene Expression Omnibus up- and downregulated genes (GEOPertUp, GEOPertDown) and the Drug Signature Database (DSigDB).

**Table S8**. Significant overlaps between genes in the “salmon” WGCNA module and transcriptomic signatures from up- and downregulated genes in the Gene Expression Omnibus (GEOPertUp and GEOPertDown, respectively), the Drug Signature Database (DSigDB) and the DrugMatrix Database.

**Table S9**. Comparison of the differentially expressed genes and WGCNA “salmon” gene lists with the results reported by Smith et al. 2016.

## References

Aarsland, D., Bronnick, K., Williams-Gray, C., Weintraub, D., Marder, K., Kulisevsky, J., Burn, D., Barone, P., Pagonabarraga, J., Allcock, L., Santangelo, G., Foltynie, T., Janvin, C., Larsen, J.P., Barker, R.A., Emre, M., 2010. Mild cognitive impairment in Parkinson disease: A multicenter pooled analysis. Neurology 75, 1062–1069.

Allen Institute for Brain Science, 2015. Allen Cell Types Database RNA-Seq Data Navig. Mouse. URL: http://celltypes.brain-map.org/rnaseq/mouse/v1-alm (accessed 1.20.20).

Armstrong, R.A., 2017. Laminar degeneration of frontal and temporal cortex in Parkinson disease dementia. Neurol. Sci. 38, 667–671.

Benito, E., Barco, A., 2015. The Neuronal Activity-Driven Transcriptome. Mol. Neurobiol. 51, 1071–1088.

Berke, J.D., Paletzki, R.F., Aronson, G.J., Hyman, S.E., Gerfen, C.R., 1998. A complex program of striatal gene expression induced by dopaminergic stimulation. J. Neurosci. 18, 5301–10.

Bezard, E., Yue, Z., Kirik, D., Spillantini, M.G., 2013. Animal models of Parkinson’s disease: Limits and relevance to neuroprotection studies. Mov. Disord. 28, 61–70.

Braak, H., Del Tredici, K., Rüb, U., De Vos, R.A.I., Jansen Steur, E.N.H., Braak, E., 2003. Staging of brain pathology related to sporadic Parkinson’s disease. Neurobiol. Aging 24, 197–211.

Buddhala, C., Loftin, S.K., Kuley, B.M., Cairns, N.J., Campbell, M.C., Perlmutter, J.S., Kotzbauer, P.T., 2015. Dopaminergic, serotonergic, and noradrenergic deficits in Parkinson disease. Ann. Clin. Transl. Neurol. 2, 949–959.

Chen, E.Y., Tan, C.M., Kou, Y., Duan, Q., Wang, Z., Meirelles, G. V., Clark, N.R., Ma’ayan, A., 2013. Enrichr: Interactive and collaborative HTML5 gene list enrichment analysis tool. BMC Bioinformatics 14.

Cools, R., Barker, R.A., Sahakian, B.J., Robbins, T.W., 2003. L-Dopa medication remediates cognitive inflexibility, but increases impulsivity in patients with Parkinson’s disease. Neuropsychologia 41, 1431–1441.

De Deurwaerdère, P., Giovanni, G. Di, Millan, M.J., 2017. Expanding the repertoire of L-DOPA’s actions: A comprehensive review of its functional neurochemistry. Prog. Neurobiol. 151, 57–100.

Duke, D.C., Moran, L.B., Kalaitzakis, M.E., Deprez, M., Dexter, D.T., Pearce, R.K.B., Graeber, M.B., 2006. Transcriptome analysis reveals link between proteasomal and mitochondrial pathways in Parkinson’s disease. Neurogenetics 7, 139–148.

Dumitriu, A., Golji, J., Labadorf, A.T., Gao, B., Beach, T.G., Myers, R.H., Longo, K.A., Latourelle, J.C., 2016. Integrative analyses of proteomics and RNA transcriptomics implicate mitochondrial processes, protein folding pathways and GWAS loci in Parkinson disease. BMC Med. Genomics 9, 5.

Dumitriu, A., Latourelle, J.C., Hadzi, T.C., Pankratz, N., Garza, D., Miller, J.P., Vance, J.M., Foroud, T., Beach, T.G., Myers, R.H., 2012. Gene expression profiles in Parkinson disease prefrontal cortex implicate FOXO1 and genes under its transcriptional regulation. PLoS Genet. 8.

Durinck, S., Moreau, Y., Kasprzyk, A., Davis, S., De Moor, B., Brazma, A., Huber, W., 2005. BioMart and Bioconductor: a powerful link between biological databases and microarray data analysis. Bioinformatics 21, 3439–40.

Ghilardi, M.F., Feigin, A.S., Battaglia, F., Silvestri, G., Mattis, P., Eidelberg, D., Di Rocco, A., 2007. L-Dopa infusion does not improve explicit sequence learning in Parkinson’s disease. Park. Relat. Disord. 13, 146–151.

González-Redondo, R., García-García, D., Clavero, P., Gasca-Salas, C., García-Eulate, R., Zubieta, J.L., Arbizu, J., Obeso, J.A., Rodríguez-Oroz, M.C., 2014. Grey matter hypometabolism and atrophy in Parkinson’s disease with cognitive impairment: A two-step process. Brain 137, 2356–2367.

Halliday, G.M., Leverenz, J.B., Schneider, J.S., Adler, C.H., 2014. The neurobiological basis of cognitive impairment in Parkinson’s disease. Mov. Disord. 29, 634–650.

Hauser, D.N., Hastings, T.G., 2013. Mitochondrial dysfunction and oxidative stress in Parkinson’s disease and monogenic parkinsonism. Neurobiol. Dis. 51, 35–42.

Heiman, M., Heilbut, A., Francardo, V., Kulicke, R., Fenster, R.J., Kolaczyk, E.D., Mesirov, J.P., Surmeier, D.J., Cenci, M.A., Greengard, P., 2014. Molecular adaptations of striatal spiny projection neurons during levodopa-induced dyskinesia. Proc. Natl. Acad. Sci. 111, 4578–4583.

Herdegen, T., Leah, J.D., 1998. Inducible and constitutive transcription factors in the mammalian nervous system: Control of gene expression by Jun, Fos and Krox, and CREB/ATF proteins. Brain Res. Rev.

Hornykiewicz, O., 1998. Biochemical aspects of Parkinson’ s disease. Neurology 51, S2–S9.

Hoss, A.G., Labadorf, A., Beach, T.G., Latourelle, J.C., Myers, R.H., 2016. microRNA Profiles in Parkinson’s Disease Prefrontal Cortex. Front. Aging Neurosci. 8, 36.

Kamińska, K., Lenda, T., Konieczny, J., Czarnecka, A., Lorenc-Koci, E., 2017. Depressive-like neurochemical and behavioral markers of Parkinson’s disease after 6-OHDA administered unilaterally to the rat medial forebrain bundle. Pharmacol. Reports 69, 985–994.

Kelly, C., de Zubicaray, G., Di Martino, A., Copland, D.A., Reiss, P.T., Klein, D.F., Castellanos, F.X., Milham, M.P., McMahon, K., 2009. L-dopa modulates functional connectivity in striatal cognitive and motor networks: a double-blind placebo-controlled study. J. Neurosci. 29, 7364–78.

Kim, D., Paggi, J.M., Park, C., Bennett, C., Salzberg, S.L., 2019. Graph-based genome alignment and genotyping with HISAT2 and HISAT-genotype. Nat. Biotechnol. 37, 907–915.

Kitano, J., Kimura, K., Yamazaki, Y., Soda, T., Shigemoto, R., Nakajima, Y., Nakanishi, S., 2002. Tamalin, a PDZ domain-containing protein, links a protein complex formation of group 1 metabotropic glutamate receptors and the guanine nucleotide exchange factor cytohesins. J. Neurosci. 22, 1280–1289.

Konradi, C., Westin, J.E., Carta, M., Eaton, M.E., Kuter, K., Dekundy, A., Lundblad, M., Cenci, M.A., 2004. Transcriptome analysis in a rat model of L-DOPA-induced dyskinesia. Neurobiol. Dis. 17, 219–236.

Kreiner, G., 2015. Compensatory mechanisms in genetic models of neurodegeneration: are the mice better than humans? Front. Cell. Neurosci. 9, 1–6.

Kuleshov, M. V., Jones, M.R., Rouillard, A.D., Fernandez, N.F., Duan, Q., Wang, Z., Koplev, S., Jenkins, S.L., Jagodnik, K.M., Lachmann, A., McDermott, M.G., Monteiro, C.D., Gundersen, G.W., Ma’ayan, A., 2016. Enrichr: a comprehensive gene set enrichment analysis web server 2016 update. Nucleic Acids Res. 44, W90–W97.

Langfelder, P., Horvath, S., 2008. WGCNA: An R package for weighted correlation network analysis. BMC Bioinformatics 9.

Lorenc-Koci, E., Czarnecka, A., Lenda, T., Kamińska, K., Konieczny, J., 2013. Molsidomine, a nitric oxide donor, modulates rotational behavior and monoamine metabolism in 6-OHDA lesioned rats treated chronically with L-DOPA. Neurochem. Int. 63, 790–804.

Martinez-Martin, P., Schapira, A.H.V., Stocchi, F., Sethi, K., Odin, P., MacPhee, G., Brown, R.G., Naidu, Y., Clayton, L., Abe, K., Tsuboi, Y., MacMahon, D., Barone, P., Rabey, M., Bonuccelli, U., Forbes, A., Breen, K., Tluk, S., Olanow, C.W., Thomas, S., Rye, D., Hand, A., Williams, A.J., Ondo, W., Chaudhuri, K.R., 2007. Prevalence of nonmotor symptoms in Parkinson’s disease in an international setting; study using nonmotor symptoms questionnaire in 545 patients. Mov. Disord. 22, 1623–1629.

Matosin, N., Fernandez-Enright, F., Lum, J.S., Andrews, J.L., Engel, M., Huang, X.F., Newell, K.A., 2014. Metabotropic glutamate receptor 5, and its trafficking molecules Norbin and Tamalin, are increased in the CA1 hippocampal region of subjects with schizophrenia. Schizophr. Res. 166, 212–218.

Mattila, P., Roytta, M., Lonnberg, P., Marjamaki, P., Helenius, H., Rinne, J.O., 2001. Choline acetyltransferase activity and striatal dopamine receptors in Parkinson’s disease in relation to cognitive impairment. Acta Neuropathol. 102, 160–166.

Mihaescu, A.S., Masellis, M., Graff-Guerrero, A., Kim, J., Criaud, M., Cho, S.S., Ghadery, C., Valli, M., Strafella, A.P., 2019. Brain degeneration in Parkinson’s disease patients with cognitive decline: a coordinate-based meta-analysis. Brain Imaging Behav. 13, 1021–1034.

Morrish, P.K., Rakshi, J.S., Bailey, D.L., Sawle, G. V., Brooks, D.J., 1998. Measuring the rate of progression and estimating the preclinical period of Parkinson’s disease with [18F] dopa PET. J. Neurol. Neurosurg. Psychiatry 64, 314–319.

O’Callaghan, C., Shine, J.M., Lewis, S.J.G., Hornberger, M., 2014. Neuropsychiatric symptoms in Parkinson’s disease: Fronto-striatal atrophy contributions. Parkinsonism Relat. Disord. 20, 867–872.

Perese, D.A., Ulman, J., Viola, J., Ewing, S.E., Bankiewicz, K.S., 1989. A 6-hydroxydopamine-induced selective parkinsonian rat model. Brain Res. 494, 285–293.

Piechota, M., Korostynski, M., Ficek, J., Tomski, A., Przewlocki, R., 2016. Seqinspector: position-based navigation through the ChIP-seq data landscape to identify gene expression regulators. BMC Bioinformatics 17, 85.

Piechota, M., Korostynski, M., Sikora, M., Golda, S., Dzbek, J., Przewlocki, R., 2012. Common transcriptional effects in the mouse striatum following chronic treatment with heroin and methamphetamine. Genes, Brain Behav. 11, 404–414.

Riedel, O., Klotsche, J., Spottke, A., Deuschl, G., Förstl, H., Henn, F., Heuser, I., Oertel, W., Reichmann, H., Riederer, P., Trenkwalder, C., Dodel, R., Wittchen, H.U., 2010. Frequency of dementia, depression, and other neuropsychiatric symptoms in 1,449 outpatients with Parkinson’s disease. J. Neurol. 257, 1073–1082.

Riekkinen, M., Kejonen, K., Ja, P., 1998. Reduction of noradrenaline impairs attention and dppamine depletion slows responses in Parkinson’s disease. Eur. J. Neurosci. 10, 1429–1435.

Riley, B.E., Gardai, S.J., Emig-Agius, D., Bessarabova, M., Ivliev, A.E., Schüle, B., Alexander, J., Wallace, W., Halliday, G.M., Langston, J.W., Braxton, S., Yednock, T., Shaler, T., Johnston, J.A., 2014. Systems-based analyses of brain regions functionally impacted in Parkinson’s disease reveals underlying causal mechanisms. PLoS One 9.

Rowe, J., Stephan, K.E., Friston, K., Frackowiak, R., Lees, A., Passingham, R., 2002. Attention to action in Parkinson’s disease: impaired effective connectivity among frontal cortical regions. Brain 125, 276–89.

Sawada, Y., Nishio, Y., Suzuki, K., Hirayama, K., Takeda, A., Hosokai, Y., Ishioka, T., Itoyama, Y., Takahashi, S., Fukuda, H., Mori, E., 2012. Attentional Set-Shifting Deficit in Parkinson’s Disease Is Associated with Prefrontal Dysfunction: An FDG-PET Study. PLoS One 7, e38498.

Schintu, N., Zhang, X., Stroth, N., Mathé, A.A., Andrén, P.E., Svenningsson, P., 2020. Non-dopaminergic Alterations in Depression-Like FSL Rats in Experimental Parkinsonism and L-DOPA Responses. Front. Pharmacol. 11, 304.

Shiner, T., Symmonds, M., Guitart-Masip, M., Fleming, S.M., Friston, K.J., Dolan, R.J., 2015. Dopamine, salience, and response set shifting in prefrontal cortex. Cereb. Cortex 25, 3629–3639.

Simioni, A.C., Dagher, A., Fellows, L.K., 2017. Effects of levodopa on corticostriatal circuits supporting working memory in Parkinson’s disease. Cortex 93, 193–205.

Simola, N., Morelli, M., Carta, A.R., 2007. The 6-Hydroxydopamine model of parkinson’s disease. Neurotox. Res. 11, 151–167.

Smith, L.M., Parr-Brownlie, L.C., Duncan, E.J., Black, M.A., Gemmell, N.J., Dearden, P.K., Reynolds, J.N.J., 2016. Striatal mRNA expression patterns underlying peak dose l-DOPA-induced dyskinesia in the 6-OHDA hemiparkinsonian rat. Neuroscience 324, 238–251.

Tasic, B., Yao, Z., Graybuck, L.T., Smith, K.A., Nguyen, T.N., Bertagnolli, D., Goldy, J., Garren, E., Economo, M.N., Viswanathan, S., Penn, O., Bakken, T., Menon, V., Miller, J., Fong, O., Hirokawa, K.E., Lathia, K., Rimorin, C., Tieu, M., Larsen, R., Casper, T., Barkan, E., Kroll, M., Parry, S., Shapovalova, N. V., Hirschstein, D., Pendergraft, J., Sullivan, H.A., Kim, T.K., Szafer, A., Dee, N., Groblewski, P., Wickersham, I., Cetin, A., Harris, J.A., Levi, B.P., Sunkin, S.M., Madisen, L., Daigle, T.L., Looger, L., Bernard, A., Phillips, J., Lein, E., Hawrylycz, M., Svoboda, K., Jones, A.R., Koch, C., Zeng, H., 2018. Shared and distinct transcriptomic cell types across neocortical areas. Nature 563, 72–78.

Trapnell, C., Williams, B.A., Pertea, G., Mortazavi, A., Kwan, G., Van Baren, M.J., Salzberg, S.L., Wold, B.J., Pachter, L., 2010. Transcript assembly and quantification by RNA-Seq reveals unannotated transcripts and isoform switching during cell differentiation. Nat. Biotechnol. 28, 511–515.

U.S. Department of Health and Human Services, 2010. National Toxicology Program. DrugMatrix. URL: https://ntp.niehs.nih.gov/data/drugmatrix/ (accessed 04.24.2020).

Ungerstedt, U., 1968. 6-Hydroxy-Dopamine Induced Degeneration of Central Monoamine Neurons. Eur. J. Pharmacol. 5, 107–110.

Vercruysse, S., Spildooren, J., Heremans, E., Wenderoth, N., Swinnen, S.P., Vandenberghe, W., Nieuwboer, A., 2014. The neural correlates of upper limb motor blocks in Parkinson’s disease and their relation to freezing of gait. Cereb. Cortex 24, 3154–66.

Walker, M.D., Dinelle, K., Kornelsen, R., Lee, A., Farrer, M.J., Stoessl, A.J., Sossi, V., 2013. Measuring dopaminergic function in the 6-OHDA-lesioned rat: A comparison of PET and microdialysis. EJNMMI Res. 3, 1–11.

Yoo, M., Shin, J., Kim, J., Ryall, K.A., Lee, K., Lee, S., Jeon, M., Kang, J., Tan, A.C., 2015. DSigDB:Drug signatures database for gene set analysis. Bioinformatics 31, 3069–3071.

Zygmunt, M., Piechota, M., Rodriguez Parkitna, J., Korostyński, M., 2019. Decoding the transcriptional programs activated by psychotropic drugs in the brain. Genes, Brain Behav. 18, e12511.

